# CellClick: an interactive platform for adjustable and accurate cell type annotation in single-cell and spatial omics data

**DOI:** 10.64898/2026.06.01.727775

**Authors:** Lei Shi, Min Dai, Yongbo Zhang, Song Wu, Meng Wang, Xiu-Jie Wang

**Affiliations:** Department of Medical Artificial Intelligence, South China Hospital, Medical School, Shenzhen University, Shenzhen, 518116, China; Institute of Genetics and Developmental Biology, Chinese Academy of Sciences, Beijing 100101, China; School of Future Technology, University of Chinese Academy of Sciences, Beijing 100049, China

**Keywords:** single-cell omics, spatial omics, cell type annotation, COSG, interactive analysis platform

## Abstract

Single-cell omics and spatial omics technologies are nowadays widely used in biological and medical research. In both single-cell and spatial omics data analysis, accurate cell type annotation is a key step for downstream analysis and scientific discoveries. However, high-quality cell annotation usually requires multiple rounds of manual analysis for result refinement, which poses great challenges to most researchers. Here, we present CellClick, an interactive platform for convenient and accurate cell type annotation in single-cell and spatial omics data. CellClick provides Data Preprocessing, Data Visualization, Cell Annotation, Annotation Validation, and Cell Reannotation modules, which facilitate automatic or user-guided cell selection and annotation. The feasibility of using CellClick to generate more accurate cell annotation results was exemplified by both scRNA-seq and spatial transcriptomics data.

## 1. Introduction

Single-cell omics and spatial omics technologies have brought unprecedented power to reveal the heterogeneity of cells within a given sample, and have promoted many important discoveries in life science and medical research. Despite their advantages, both single-cell and spatial omics technologies rely on comprehensive data analysis process to obtain valuable discoveries, which have created hardships for users. Cell type annotation is a basic and essential step in both single-cell and spatial omics data analysis, which determines the accuracy of downstream results ^[1-5]^. Although many methods have been developed, the cell annotation step is still error-prone and difficult for most researchers ^[5-11]^.

In practice, software-based automatic cell annotation and expert-based manual cell annotation are two mainstream approaches for cell type identification. As the software-based automatic annotation methods mainly rely on pre-defined cell marker genes or the analysis results of existing single-cell omics data, they are prone to inherit mistakes and have difficulties to identify novel cell types ^[12, 13]^. Therefore, manual cell annotation is necessary and sometimes needs to be performed multiple rounds to achieve accurate cell type identification results. Manual cell annotation methods usually require researchers to evaluate the reliability of cell clustering results, then redefine the marker genes and annotations for some cell groups. Such comprehensive process is unattainable for most researchers when the analysis software requires programmable-skill-based interactions.

To facilitate user-friendly cell annotation, efforts have been made to develop interactive analysis platforms to eliminate the requirement for programming skills from users ^[14-17]^. However, the currently available single-cell or spatial omics data analysis platforms either offer limited functions or do not support multiple rounds of cell type identification analysis, which makes it difficult to select and reannotate the inappropriately or suspiciously annotated cells. Here we present CellClick, an easy-to-use interactive software for single-cell and spatial omics data analysis. CellClick provides comprehensive data analysis tools and interactive cell annotation/reannotation methods, which offers an easy-to-use tool for inexperienced users to perform cell annotation on single-cell or spatial omics data.

## 2. Methods

### 2.1 Single-Cell Omics Data Collection and Preprocessing

#### 3k PBMCs dataset

The 3k PBMCs dataset, a human peripheral blood single-cell RNA-seq dataset, was downloaded from the 10x Genomics website (https://support.10xgenomics.com/single-cell-gene-expression/datasets/1.1.0/pbmc3k) and used to illustrate the main functions of CellClick. The preprocessing of 3k PBMCs was performed using the Preprocessing module of CellClick, which included quality control (QC), normalization, highly variable gene (HVG) detection, dimension reduction, and cell clustering functions. Low-quality cells were filtered out if a cell had any of the following features: 1) with fewer than 400 total UMI counts or more than 7,600 total UMI counts; 2) with fewer than 200 detected genes or more than 2,000 detected genes; 3) with 5% or more UMI counts derived from mitochondrial genomes. During normalization, the count matrix was normalized to 10,000 per cell, followed by log-transformation. For HVG detection, the top 2,000 highly variable genes were identified using the dispersion-based method ^[18]^. For dimensionality reduction, UMAP was applied to capture both the local and global structures, and cells were clustered using the Leiden algorithm ^[9]^.

#### 10k PBMCs dataset

The 10k PBMCs dataset, a human peripheral blood single-cell RNA-seq dataset, was downloaded from the 10x Genomics website (https://www.10xgenomics.com/cn/datasets/10-k-pbm-cs-from-a-healthy-donor-v-3-chemistry-3-standard-3-0-0) and used to demonstrate the ability of CellClick in generating more accurate cell type annotation results for preprocessed single-cell RNA-seq dataset. The 10k PBMCs dataset was first treated with Scanpy v1.9.3 [19], scrublet v0.2.3 [20], and scikit-learn v1.0.2 [21] for QC analysis. Low-quality cells were filtered out if a cell had any of the following features: 1) with fewer than 500 total UMI counts or more than 25,000 total UMI counts; 2) with fewer than 1,000 detected genes or more than 4,500 detected genes; 3) with 20% or more UMI counts derived from mitochondrial genomes; 4) with no less than 0.2 doublet score calculated by Scrublet [20] with default parameters; 5) with abnormal values in any of the above attributes detected by the IQR (Interquartile Range) method. After QC, the preprocessed 10k PBMCs dataset was uploaded into CellClick for downstream analysis, and cells of the 10k PBMCs dataset were clustered into 8 cell clusters using the Leiden algorithm [9]. For cell type annotation, the reference marker genes from CellMarker 2.0 were used as the input of the Cell Identification function. In addition, the human PBMC cell atlas from the ToppCell database [22] was used as the reference data for 10k PBMCs to validate the accuracy of cell annotation results with the Reference Comparison function.

#### Mouse E2S1 dataset

The Mouse E2S1 dataset was downloaded from mouse organogenesis spatiotemporal transcriptomic atlas (https://ftp.cngb.org/pub/SciRAID/stomics/STDS0000058/stomics/E10.5_E2S1.MOSTA.h5ad) and used to demonstrate the advantage of CellClick in analyzing spatial transcriptomic data. To better illustrate the results, cells do not belong to clusters “Brain”, “Dorsal root ganglion”, “Jaw and tooth”, and “Mucosal epithelium”, according to the original annotations, were reannotated as cluster “Others” in this work.

### 2.2 Development of CellClick

CellClick was developed in Python 3.10. The user interface of CellClick was built using Plotly/Dash v2.1.0 framework, which enables the rapid development of interactive data applications. The back-end of CellClick is supported by several Python libraries, including Scanpy v1.8.2 [19], COSG v1.0.1 [23], SciPy v1.7.3 [24], NumPy 1.21.6 [25], and pandas 1.3.5 [26]. In CellClick, the single-cell omics data are stored as Annotated Data (AnnData) objects which leverage the HDF5 library for efficient data management [27]. To support more accurate cell type annotation, CellClick provided five modules, namely Data Preprocessing, Data Visualization, Cell Annotation, Annotation Validation, and Cell Reannotation modules. Besides the command-line execution, CellClick also provides a web-based interface and can be executed in a Jupyter environment.

#### Data Preprocessing Module

CellClick accepts single-cell omics data in the h5ad format and enables a standard preprocessing workflow with the built-in functions of Scanpy ^[19]^ for unpreprocessed datasets, including the QC, data normalization, HVG detection, and dimension reduction functions (Figure 1A). Dash components are employed to generate interactive data forms and graphs in this module.

**Figure 1.**
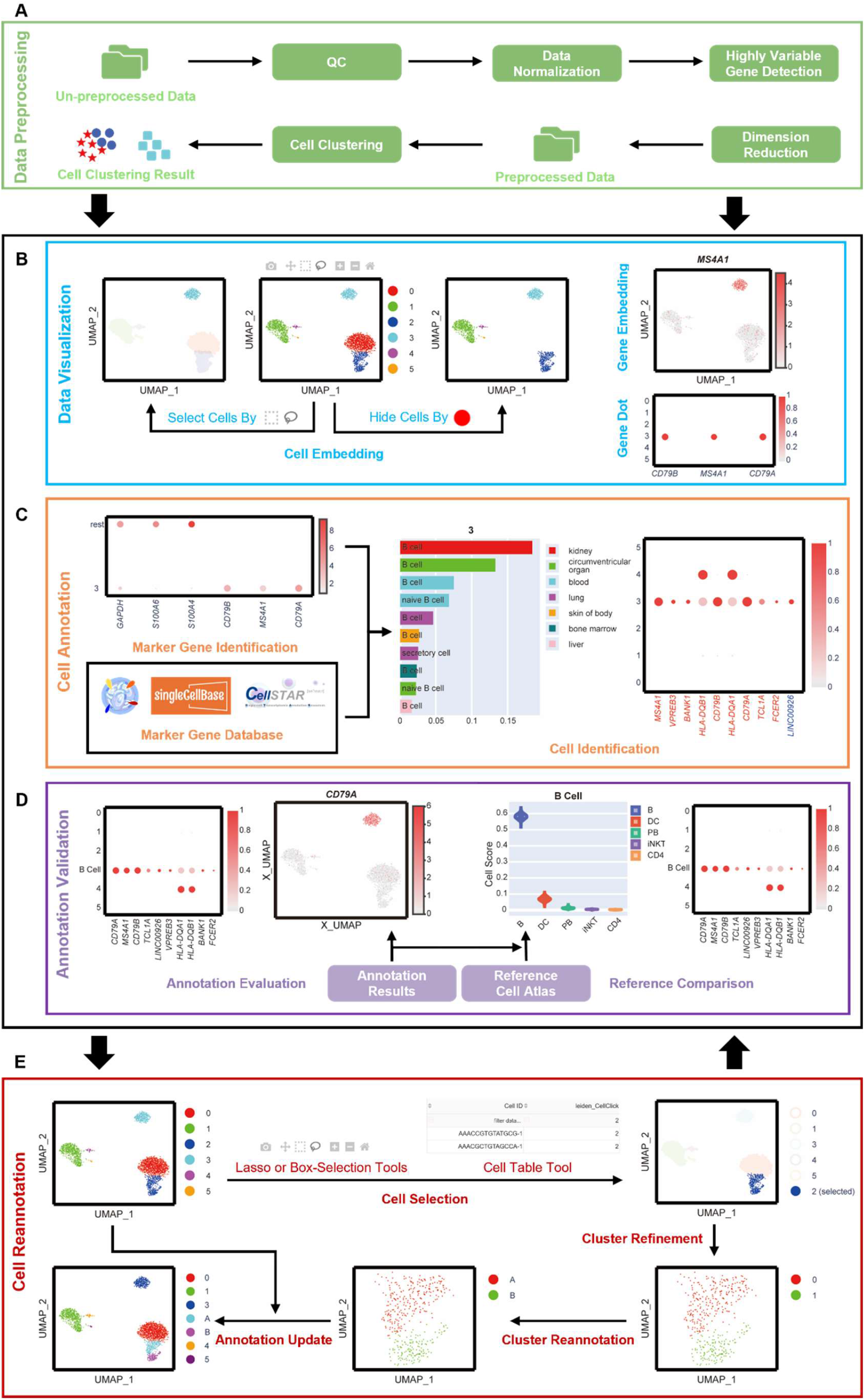
The workflow and main functions of CellClick. **(A)** Workflow and functions of the Data Preprocessing module. **(B)** Workflow and functions of the Data Visualization module. **(C)** Workflow and functions of the Cell Annotation module. In the dot plot of the Cell Identification function, the names of known reference marker genes reported by COSG are shown in red, and the names of marker genes newly identified by COSG are shown in blue. **(D)** Workflow and functions of the Annotation Validation module. The dot plot produced by the Reference Comparison function shows the expression pattern of marker genes identified for the target cluster, sorted by their COSG scores. **(E)** Workflow and functions of the Cell Reannotation module. Panels **(B)** to **(E)** represent the analysis results of the 3k PBMCs dataset.

#### Data Visualization Module

To enable interactive data analysis, CellClick provides diverse methods for data visualization and analysis. All graph displays are generated by Dash components, and the callback functions of Dash package are used for real-time result display and event-driven result update. For example, users can click the legend to hide or redisplay a selected cell cluster, thereby to support user-defined data selection and visualization. To support parallel visualization of analysis results, CellClick employs Dash package to generate multiple canvases (up to 6 by default). All graph display functions are built upon a template-driven rendering framework that enables interactive data visualization.

#### Cell Annotation Module

In the Cell Annotation module, the Marker Gene Identification function is achieved via COSG ^[23]^, and the Cell Identification function is achieved with the following method.

The reference marker genes were downloaded from CellMarker 2.0 (http://bio-bigdata.hrbmu.edu.cn/CellMarker/CellMarker_download.html) ^[28]^, singleCellBase (http://cloud.capitalbiotech.com/SingleCellBase/download.jsp) ^[29]^, and CellStar (https://cellstar.idrblab.net/download) ^[30]^, and prestored in CellClick, which included the gene symbol, expressed cell type, and associated publication information for each marker gene. To evaluate whether a candidate gene (*g*) is the marker gene of a reference cell type (*C*^*r*^), we calculated 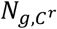 (the number of publications referring *g* as the marker gene of *C*^*r*^), *N*_*g*_ (the number of publications referring *g* as a marker gene of any cell type *C*), and 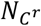 (the number of publications mentioning any known marker genes for *C*^*r*^), and defined the marker gene confidence score (MGC) of *g* for *C*^*r*^ with the equation:

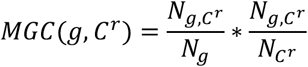

To achieve more reliable results, only gene-cell type pairs with 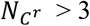 were included.

As some marker genes identified by COSG may be new and not included in previous publications, we defined the overlapping marker score (OMS) to calculate the overlapping status between COSG identified marker genes and the reference markers for any given cell type using the following equation:

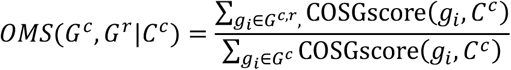

where *G*^*c,r*^ represents the overlapped marker genes between 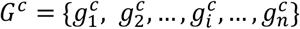 (COSG identified marker genes for a given cell cluster *C*^*c*^) and 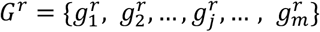 (the reference marker genes for any reference cell type *C*^*r*^ recorded in CellMarker 2.0 ^[28]^, singleCellBase ^[29]^, or CellStar ^[30]^). Besides, COSGscore(*g*_*i*_, *C*^*c*^) represents the expression specificity score of gene *g*_*i*_ in *C*^*c*^ calculated by COSG ^[23]^.

With *OMS*(*G*^*c*^, *G*^*r*^|*C*^*c*^) and *MGC*(*g, C*^*r*^), the Cell Identification function defines the confidence of annotating *C*^*c*^ as *C*^*r*^ by the equation:

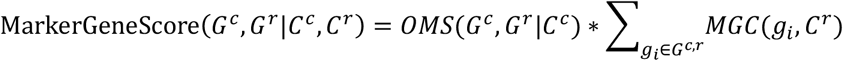

#### Annotation Validation Module

The Annotation Validation module includes the Annotation Evaluation and Reference Comparison functions. The Annotation Evaluation function was developed using Dash components. The Reference Comparison function compares the similarity of marker genes identified in the analyzed dataset with the marker genes identified from the reference cell atlas (Table S1), both by the COSG method, via the following functions.

Given a set of marker genes 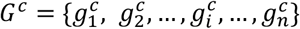 for a defined cell type *C*^*c*^, and the marker gene set 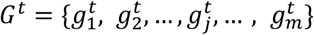 obtained via the COSG method for the reference cell type *C*^*r*^, the Reference Comparison function calculates CellScore(*c*_*i*_|*C*^*c*^, *C*^*r*^, *G*^*c*^, *G*^*t*^) to estimate the reliability of annotating a cell *c*_*i*_ from *C*^*c*^ as *C*^*r*^ using the following equation:

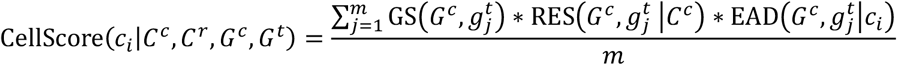

In the above equation, GS represents the similarity between *G*^*c*^ and 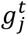 based on cosine similarity, which is calculated by:

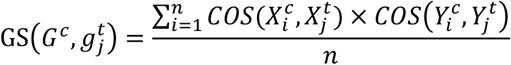

where 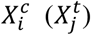 represents the expression level of 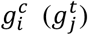 in all cells, 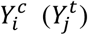 represents the mean expression level of 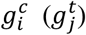 within each cell cluster, and *COS* denotes the cosine function.

RES represents the relative expression specificity of 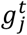 as compared with *G*^*c*^ in a given cell cluster *C*^*c*^, and is calculated by:

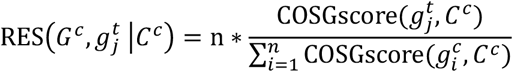

where COSGscore(*g*_*i*_, *C*^*c*^) is generated by COSG ^[23]^ and represents the expression specificity of gene *g*_*i*_ in a given cell cluster *C*^*c*^.

EAD represents the expression abundance difference of 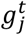 compared with *G*^*c*^ in *c*_*i*_ by a sigmoid-like function, and is calculated by:

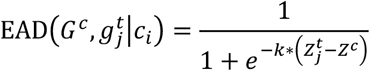

where 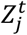 and *Z*^*c*^ represent the expression level of 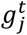 in *c*_*i*_ and the mean expression level of *G*^*c*^ in *c*_*i*_, respectively. And *k* (equals to 0.5 by default) denotes the scale factor of expression abundance difference.

Note that 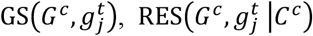, and 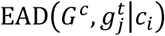 are all treated as 1 when 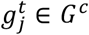. Generally, a higher cell score distribution means the expression patterns of genes in *G*^*t*^ and *G*^*c*^ are more similar, which always demonstrates a greater reliability to annotate *C*^*c*^ as *C*^*t*^.

#### Reannotation Module

The Reannotation module develops an iterative refinement framework for correcting annotation errors in single-cell omics datasets, including Cell Selection, Cluster Refinement, Cluster Reannotation, and Annotation Update functions, which were all developed using Dash and Scanpy ^[19]^.

### 2.3 Code availability

CellClick is available at https://omicslab.genetics.ac.cn/download/CellClick.zip and https://github.com/Omics-Shi4712/CellClick.

## 3. Results

### 3.1 Overview of CellClick functions

CellClick is a web-based data analysis platform with interactive functions specialized for cell type identification in single-cell omics or spatial omics data (Figure 1). CellClick takes different types of single-cell or spatial omics sequencing data (either unpreprocessed or preprocessed) with the Scanpy-supported formats (e.g., the h5ad format) as inputs ^[19]^. The major feature of CellClick is the ability to provide multiple interactive functions for users to preprocess, visualize, annotate, evaluate, re-annotate or re-cluster cells. It also allows parallel comparison of the analysis results for different genes, cell clusters, or datasets, which is currently absent in most available software. CellClick provides the above-mentioned functions in five modules, namely Data Preprocessing, Data Visualization, Cell Annotation, Annotation Validation, and Cell Reannotation modules (Figure 1).

#### Data Preprocessing

When an unpreprocessed dataset is uploaded, users can perform QC analysis and invoke the general analysis functions of Scanpy ^[19]^ (e.g., data normalization, HVG detection, dimension reduction, and cell clustering) through CellClick (Figure 1A). In CellClick, low-quality cells can be filtered based on common covariates during the QC step (e.g., UMI counts, gene counts, or percentage of mitochondrial gene counts). CellClick also provides a user-friendly interface for data normalization, HVG detection, dimension reduction, and cell clustering. When a preprocessed dataset is uploaded, users can skip the majority of data preprocessing steps and start with the Cell Clustering function or select the targeted cell clustering results in the uploaded annotation files for downstream analysis (Figure 1A).

#### Data Visualization

CellClick supports interactive functions for users to visualize cell embedding, gene embedding, and gene dot graph results (Figure 1B). The Cell Embedding function empowers direct visualization of cell features in low-dimensional space, which is essential for cell type identification (Figure 1B). To ensure accurate cell type annotation results, users often need to select target cell groups in the embedding map for refined marker gene identification or cell type reannotation. To meet such needs, CellClick allows users to select cells in the embedding results via the box-selection or lasso-selection tools, then take the selected cells for various downstream analysis (Figure 1B). By clicking on the legend icon of each cell cluster, the cell cluster can be hidden or redisplayed, thus enabling users to visualize and select target cell sets more accurately (Figure 1B). The Gene Embedding function displays the expression profile of any selected gene in the embedding space, which can be used to assess the expression specificity of candidate marker genes (Figure 1B). Besides the gene embedding function, users can also review the expression profiles of multiple genes via the Gene Dot function (Figure 1B). One unique feature of CellClick is its multi-canvas function, which allows users to compare multiple visualization results (up to 6 by default) in a parallel manner, such as generating embedding or dot plot profiles for multiple genes (Figure S1A, S1B, and S1C online). The multi-canvas function also supports parallel visualization of different datasets, thus enabling cross-experiments, cross-organs, and cross-species comparisons (Figure S1D and S1E online).

#### Cell Annotation

The Cell Annotation module provides Marker Gene Identification and Cell Identification functions for cell type annotation (Figure 1C). Marker Gene Identification is a basic and crucial step for single-cell or spatial omics sequencing data analysis. To facilitate fast and accurate marker gene identification, CellClick has implemented COSG ^[23]^, a cosine similarity based cell marker gene identification method, to identify marker genes for a selected cell cluster (Figure 1C). The Cell Identification function allows users to compare the marker gene identification results with the reference marker genes curated from previous publications. The reference marker gene lists provided by CellClick currently include the annotated marker genes from CellMarker 2.0 ^[28]^, singleCellBase ^[29]^, and CellStar ^[30]^ websites, and will be periodically updated upon the release of new cell marker resources (Methods). For a group of candidate marker genes selected for any cell cluster, the Cell Identification method compares these marker genes with the reference marker genes of each cell type, and returns the marker gene score (the reliability of defining a cell type using the candidate marker genes) in the top-ranked candidate cell types (top 10 by default) (Figure 1C). At the same time, a gene dot plot will also be generated for users to evaluate the expression specificity of the candidate marker genes in each candidate cell type (Figure 1C). In the gene dot plot, candidate marker genes predicted by COSG will be classified into two categories, previously reported reference markers and novel markers, which will be shown in different colors (Figure 1C). Based on the above information, users can easily annotate known cell types or identify novel cell types.

#### Annotation Validation

The Annotation Validation module includes Annotation Evaluation and Reference Comparison functions, which allows users to evaluate the cell type annotation results (Figure 1D). The Annotation Evaluation function invokes Gene Dot and Gene Embedding functions to display the expression profiles of identified marker genes for a chosen cell type (Figure 1D). In addition, users can also explore the expression profiles of other reference marker genes through this function, thus evaluating the expression specificity of the identified marker genes and the reliability of the cell type annotation results. The Reference Comparison function enables users to compare cell type annotation results with a reference cell atlas, which could be a single-cell atlas prestored in CellClick or uploaded by the user. Briefly, the Reference Comparison function compares the expression profile similarity of COSG-identified marker genes between the user annotated cell type and the recorded cell type in the reference cell atlas using cell score (the reliability of defining a cell as the labeled cell type using the corresponding known reference marker genes), and displays the cell score distributions of the top-ranked reference cell types in violin plots (top 5 by default) (Methods, Figure 1D). Similar to the Cell Identification function, the Reference Comparison function also generates a dot plot to show the expression of reference marker genes in user-annotated cell types (Figure 1D). As there are many confounding factors which could influence the cell type annotation results (e.g., inappropriate grouping of cell clusters, non-specific expression of reference marker genes), the Annotation Validation functions may need to be applied multiple times to ensure high-quality cell type annotation.

#### Cell Reannotation

The Cell Reannotation module incorporates Cell Selection, Cluster Refinement, Cluster Reannotation, and Annotation Update functions, which facilitates selecting target cells for refined cell annotation (Figure 1E). As a high-quality cell annotation result usually requires multiple rounds of manual examination and correction, users often need to precisely select inappropriately or suspiciously annotated cells after the Annotation Validation step for more accurate reannotation. To meet these demands, the Cell Selection function not only allows users to select target cells from the cell embedding map using the box-selection or lasso-selection tools, same as described in the Data Visualization module, but also enables more accurate cell selection by user-defined cell attributes (e.g., expression of certain genes) via the Cell Table tool (Figure 1E and S2 online). The box-selection, lasso-selection, and Cell Table tools are capable of capturing cells with different features, and users can combine the selection results of these approaches using the Fix Selection function to facilitate more accurate analysis (Figure S2 online). The Cluster Refinement function enables users to redo cell clustering on the selected cells via the Dimension Reduction or Cell Clustering functions (Figure 1E). To avoid over-clustering, the Cluster Refinement step can be skipped when a small group of homogeneous cells is selected or when the users are satisfied with the annotation results. Next, users need to run the Cluster Reannotation function to assign annotations to the selected cells, then use the Annotation Update function to combine the newly annotated cell clusters with the previously annotated ones (Figure 1E). After one or more rounds of Cell Reannotation, users are likely to obtain high-quality cell type identification results.

### 3.2 Application of CellClick in single-cell RNA-seq data

To facilitate the application of CellClick, we used the bench-marking human peripheral blood single-cell RNA-seq dataset (10k PBMCs) from the 10x Genomics company website to demonstrate the advantages of CellClick in generating more accurate cell type annotation results via a convenient online interactive manner (Figure 2A). After quality control and data preprocessing, a total of 8,711 cells from the 10k PBMCs dataset were retained and grouped into 8 cell clusters by the Leiden algorithm ^[9]^ (Figure 2A). We then employed the Cell Identification function of CellClick to perform cell cluster annotation using reference marker genes derived from the CellMarker 2.0 dataset ^[28]^ (Figure S3 online). The annotation results identified various common cell types in human peripheral blood cells, such as B cells, T cells, and monocytes (Figure S3 online). However, the above automated annotation process defined Cluster “0” as “monocyte” and defined Cluster “6” as “CD14-positive, CD16-positive monocyte”, resulting in the co-existence of two “monocyte” clusters without clear distinction from each other (Figure 2B). Such low-quality or confusing cell type annotation is a common problem associated with many automated cell annotation processes. With CellClick, users can easily redefine these cell types with the assistance of Gene Dot function or other functions. In this case, the marker gene dot plot showed that Cluster “0” specifically expresses *CD14* (a marker gene of CD14^+^ monocytes), whereas Cluster “6” specifically expresses *HES4* (a marker gene of CD16^+^ monocytes) ^[31]^ (Figure 2C). Therefore, we reannotated Cluster “0” and Cluster “6” as “CD14^+^ Monocyte” and “CD16^+^ Monocyte”, respectively (Figure 2D).

**Figure 2.**
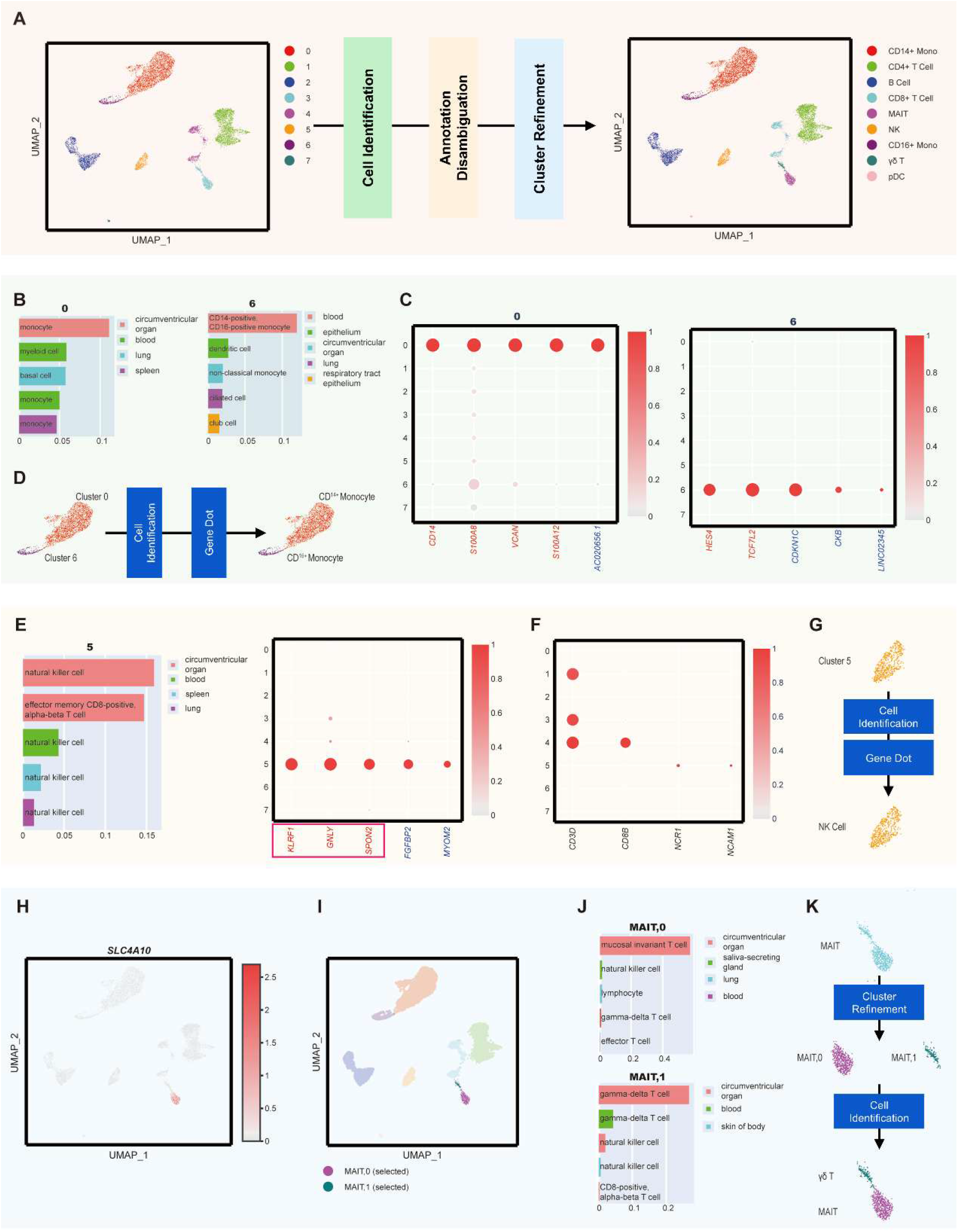
Capability of CellClick in generating more accurate cell type annotations for single-cell RNA-seq data. **(A)** The schematic diagram of the cell identification workflow for the 10k PBMCs dataset using CellClick. CD14+ Mono, CD14^+^ Monocytes; MAIT, mucosal-associated invariant T cells; NK, natural killer cells; CD16+ Mono, CD16^+^ Monocytes; pDC, plasmacytoid dendritic cells. **(B)** Reference marker gene-based marker gene scores for Cluster “0” and Cluster “6”. **(C)** Expression patterns of CellClick suggested marker genes for Cluster “0” and Cluster “6”. **(D)** Annotation results for Cluster “0” and Cluster “6” using CellClick. **(E)** Reference marker gene-based marker gene scores for Cluster “5”. Reference marker genes with ambiguous cell type identities are highlighted with red box in the dot plot. **(F)** Expression patterns of *CD3D* (the marker gene for T cells), *CD8B* (the marker gene for CD8^+^ T cells), *NCR1* (the marker gene for NK cells), and *NCAM1* (the marker gene for NK cells) across all cell clusters. **(G)** Annotation results for Cluster “5” using CellClick. **(H)** UMAP embedding showing cells expressing *SLC4A10* in the 10k PBMCs dataset. **(I)** UMAP embedding of cell selection results for Cluster “MAIT” using the Cluster Refinement function. **(J)** Newly identified marker genes for temporary Cluster “MAIT,0” and Cluster “MAIT,1” after rerunning the Cell Identification function. **(K)** Reannotation results for Cluster “MAIT” using CellClick. The color rule for gene names in the dot plots of panels **(C)** and **(E)** is the same as described in Figure 1 **(C)**.

Another common problem associated with automated cell annotation is the ambiguity of marker genes in the reference marker gene database. For example, in the CellMarker 2.0 dataset, 25.76% (1,379 of 5,354) marker genes are assigned to more than one cell type in human, and the most extreme examples are *CD4* and *PTPRC*, which are defined as the marker genes for 28 cell types. In the above mentioned automated Cell Identification results of the 10k PBMCs dataset, several identified marker genes for Cluster “5” (e.g., *KLRF1, GNLY*, and *SPON2*) were reported as marker genes for both effector memory CD8^+^ T cells and NK cells in the CellMarker 2.0 dataset ^[32-34]^, therefore users would be unable to assign a definite cell type for Cluster “5” according to the marker gene scores (Figure 2E). To generate a more accurate annotation for Cluster “5”, we used the Gene Dot function of CellClick to visualize the expression of *CD3D* (a canonical marker gene for T cells) ^[35, 36]^, *CD8B* (a canonical marker gene for CD8^+^ T cells) ^[37]^, *NCR1* (a gene encoding NK cell-specific surface molecule NKp46) ^[38]^, and *NCAM1* (a gene encoding NK cell-specific surface molecule CD56) ^[39]^ among all cell clusters (Figure 2F). The results showed that both *NCR1* and *NCAM1* were specifically expressed in Cluster “5” cells, but *CD3D* and *CD8B* were not, therefore we reannotated Cluster “5” as NK cells (Figure 2G).

It often happens that only a subset of cells within a certain cell cluster has similar gene expression patterns, but the others do not, indicating the necessity of dividing such cell cluster into more clusters. For example, in the automated 10k PBMCs annotation results, Cluster “MAIT” exhibited a tadpole-shape, and the identified marker gene *SLC4A10* for Cluster “MAIT” was only highly expressed in cells within the head region of the tadpole-shaped cluster, indicating there might be multiple cell types within Cluster “MAIT” (Figure 2H). To verify such hypothesis, we temporarily assigned the head and tail regions of the tadpole-shaped Cluster “MAIT” as Cluster “MAIT,0” and Cluster “MAIT,1”, with the Cluster Refinement function (Figure 2I), then reran the Cell Identification function for these cell clusters. Such refined cell identification process revealed marker genes of MAIT cell for Cluster “MAIT,0” and marker genes of gamma-delta (γδ) T cell for Cluster “MAIT,1”, respectively (Figure 2J). Therefore, we reannotated Cluster “MAIT” as two clusters, namely Cluster “MAIT” and Cluster “γδ T” (Figure 2K).

To evaluate the accuracy of the above cell reannotation results, we invoked the Reference Comparison function within the Annotation Validation module of CellClick, and compared the above refined cell identification results of the 10k PBMCs dataset with the PBMCs reference single-cell dataset in the ToppCell database ^[22]^. High confident cell score was obtained for the newly annotated “MAIT” Cluster (Figure S4A online) and “γδ T” Cluster (Figure S4B online). The dot plot further showed distinct marker gene expression profiles for Cluster “MAIT” and Cluster “γδ T” (Figure S4C online), respectively, demonstrating the reliability and accuracy of cell identification results generated by CellClick.

### 3.3 Application of CellClick in spatial transcriptomics data

Spatial omics techniques are capable of capturing both the molecular features and *in vivo* locations of cells, however, due to the irregular distribution of cells within tissues or organs, selecting target cell groups for accurate annotation remains challenging ^[40, 41]^. To demonstrate the convenient functions of CellClick for interactive and precise cell population selection, we collected and processed the published Mouse E2S1 dataset, a spatial transcriptome dataset of mouse E10.5 embryos, and focused on the sub-group classification of head mesenchyme cells.

The Mouse E2S1 dataset contained 8,494 cells which were divided into 18 cell clusters by the original publication ^[42]^. Among these cell clusters, head mesenchyme cells were grouped into one cluster with complicated spatial distributions (Figure 3A). Previous studies have shown that the developmental fates of head mesenchyme cells were associated with their locations. For example, supraorbital mesenchyme (SOM), lying apical to the eye, contributed to cranial bone formation, whereas the early migrating mesenchyme located above the SOM differentiated into sutures or soft tissue layers ^[43]^. In addition, cell origin analysis of the Mouse E2S1 dataset revealed that the head mesenchyme cells on the top of “Jaw and tooth” cells were primarily derived from mesoderm, whereas the head mesenchyme cells surrounded by “Mucosal epithelium” cells and “Dorsal root ganglion” cells were mainly originated from ectoderm (Figure 3B). Such distinct distribution may suggest differences in cell functions and cell fates, but clustering and analyzing these cells based on spatial information remains challenging using existing analysis platforms.

**Figure 3.**
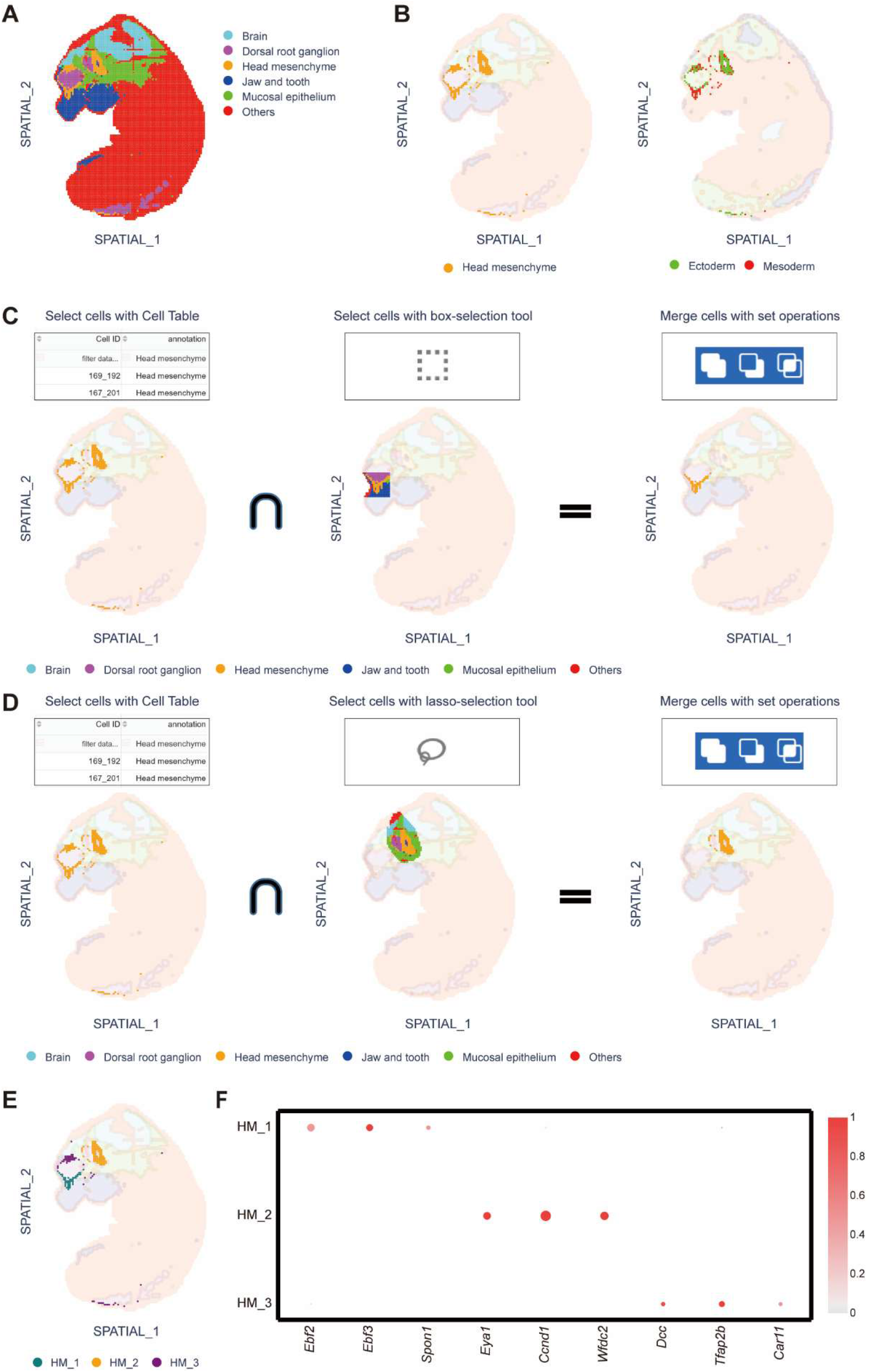
Capability of CellClick in generating more accurate cell type annotations for spatial transcriptomics data. **(A)** Spatial distribution of cell types in the Mouse E2S1 dataset with original annotation. **(B)** Spatial distribution of cells annotated by cell type or germ layer in the “Head mesenchyme” Cluster. **(C)** Specific selection of the head mesenchyme cells on the top of Cluster “Jaw and tooth” using the Cell Selection function of CellClick. **(D)** Specific selection of the head mesenchyme cells surrounded by the cells in the “Mucosal epithelium” Cluster and “Dorsal root ganglion” Cluster. **(E)** Spatial distribution of newly annotated cell Clusters “HM_1”, “HM_2”, and “HM_3”. **(F)** Expression patterns of the top 3 marker genes identified for Clusters “HM_1”, “HM_2” and “HM_3”. In Panels **(A), (C)** and **(D)**, the cluster “Others” represents all other cell clusters annotated by the original publication.

The versatile cell selection functions of CellClick can effectively solve this problem without the requirement of programming. To select cells with complicated distribution patterns, users can either apply the box-selection or lasso-selection tools, or select certain features using the Cell Table function, or use the combination of all these tools. For instance, to further annotate cells within the Cluster “Head mesenchyme”, we first isolated the Cluster “Head mesenchyme” via the Cell Table function (Figure 3C), then used the box-selection tool to select the mesoderm derived head mesenchyme cells located right above the “Jaw and tooth” cell cluster (Figure 3C). CellClick provides a merge function for users to obtain the intersections of different cell selection methods, therefore users can merge the cell selection results of the Cell Table function and the box-selection function to obtain mesoderm derived “Head mesenchyme” cells located right above the “Jaw and tooth” cell cluster (Figure 3C). By repeating the similar procedure on other “Head mesenchyme” cells, we divided the “Head mesenchyme” cells into 3 subclusters (namely HM_1, HM_2, and HM_3), representing different spatial distributions and cell origins (Figure 3C, 3D and 3E). Subsequent marker gene identification process revealed *Ebf2* and *Spon1* as marker genes for Cluster “HM_1”, indicating a role of “HM_1” cells in regulating cranial bone formation ^[44, 45]^ (Figure 3F). Therefore, we reannotated Cluster “HM_1” as SOM. The marker gene *Eya1* for Cluster “HM_2” has been reported to be required for the normal development of neural crest cells at the post-migratory stage ^[46]^, and the marker gene *Tfap2b* for Cluster “HM_3” functions to ensure proper neural crest lineage differentiation ^[47]^. These results suggested Cluster “HM_2” (*Eya1*^*+*^ HM) and Cluster “HM_3” (*Tfap2b*^*+*^ HM) as two cell subtypes of early migrating mesenchyme derived from neural crest cells, and also demonstrated the feasibility of using CellClick to improve cell type annotations for spatial transcriptomics data.

## 4. Discussion

Cell annotation is the basic yet essential step in single-cell and spatial omics sequencing data analysis, which determines the quality and reliability of downstream analysis. Yet more currently available cell annotation tools are challenging for unexperienced researchers due to their complicated operation procedures and requirements for programming. Convenient and programming-free cell annotation tools are therefore in great need.

In recent years, several interactive single-cell data analysis software packages have been developed with the aim to eliminate programming requirements from the user side ^[14-17, 48-50]^. However, these tools only provide basic cell type identification workflow and lack the result validation functions. Moreover, none of these tools support multiple rounds of cell reannotation, which is a critical step for generating accurate cell annotation results (e.g., the identification of rare cell subtypes). To address this challenge, CellClick provides the Annotation Validation and Cell Reannotation modules, which enable efficient detection and correction of inappropriately or suspiciously annotated cell types. The Annotation Validation module visualizes the expression patterns of marker genes from both user selected cells and the reference data, which facilitates the identification of inappropriately or suspiciously annotated cells. The Cell Reannotation module enables multi-round iterative optimizations of annotation results by providing versatile cell selection and annotation methods, which greatly facilitated more accurate cell type annotation.

After local installation, CellClick runs as a web-based interface with no programming requirement for common users. It also supports command-line execution for data analysis experts. Based on Dash, a framework specifically designed for interactive analytical web applications and data visualization, CellClick inherently provides an interactive framework to reveal cellular function and perform robust cell identification analysis. CellClick can be executed under the Jupyter environment, which enables seamless integration with other analysis workflows, such as the cell communication analysis using CellPhoneDB ^[51]^ and Squidpy ^[52]^. Such multi-mode design and efficient workflow would broaden the applicability of CellClick in diverse single-cell and spatial omics data analysis pipelines.

## Acknowledgments

This work was supported by the Strategic Priority Research Program of Chinese Academy of Sciences (XDA0460203) to M.W., Beijing Municipal Science & Technology Commission (Z231100007223010) and Chinese Academy of Sciences for Young Scientists in Basic Research Project (YSBR-073) to X.-J.W.

## Author contributions

X.W. supervised the study. L.S. and M.D. developed the software and built the computational tools. L.S. and Y.Z. performed the analysis, L.S., Y.Z., S.W., and M.W. contributed to the software function improvement. L.S. and X.W. wrote the manuscript. All authors read and approved the final manuscript.

