## Supplemental Figure for "CellClick: an interactive platform for adjustable and accurate cell type annotation in single-cell and spatial omics data"

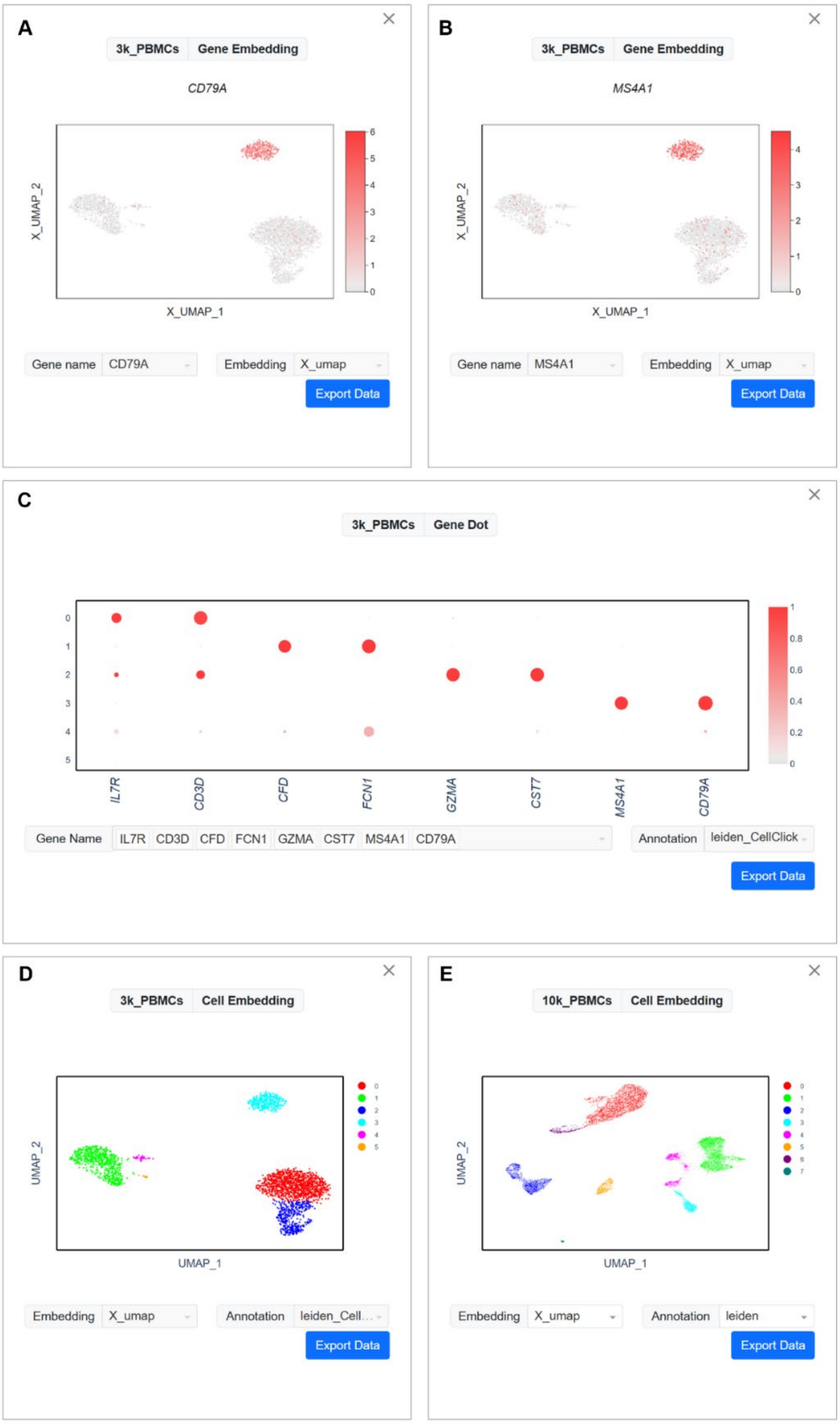


**Supplementary Figure 1. Illustration of the multi-canvas function of CellClick. (A)** UMAP embedding showing cells expressing *CD79A* in the 3k PBMCs dataset. **(B)** UMAP embedding showing cells expressing *MS4A1* in the 3k PBMCs dataset. (**C**) Dot plot showing the expression profiles of genes *IL7R*, *CD3D*, *CFD*, *FCN1*, *GZMA*, *CST7*, *MS4A1*, and *CD79A* across all cell clusters. **(D)** UMAP embedding showing the cell clustering results of the 3k PBMCs dataset. **(E)** UMAP embedding showing the cell type annotation results of the 10k PBMCs dataset.

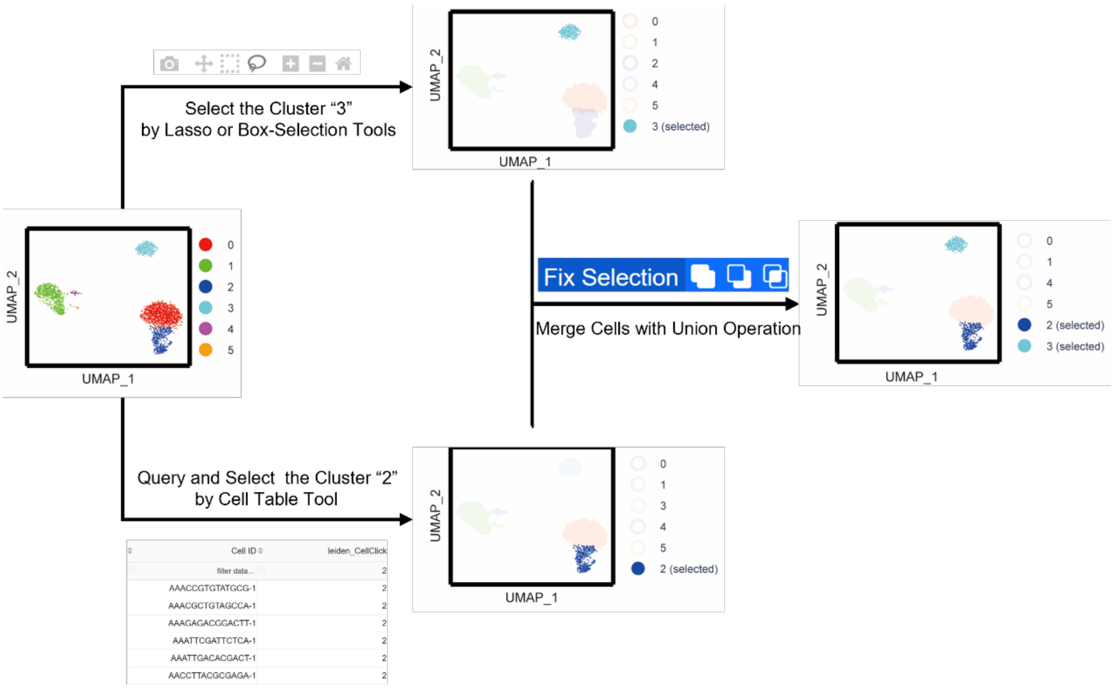


**Supplementary Figure 2. Illustration of the Cell Selection function of CellClick.** Well-separated cell clusters in UMAP (e.g., Cluster “3”) can be selected using the lasso or box-selection tools. Cell clusters adjacent to other clusters (e.g., Cluster “2”) can be selected through the Cell Table tool. To include cells from both clusters, the Fix Selection function can be applied to perform a union operation.

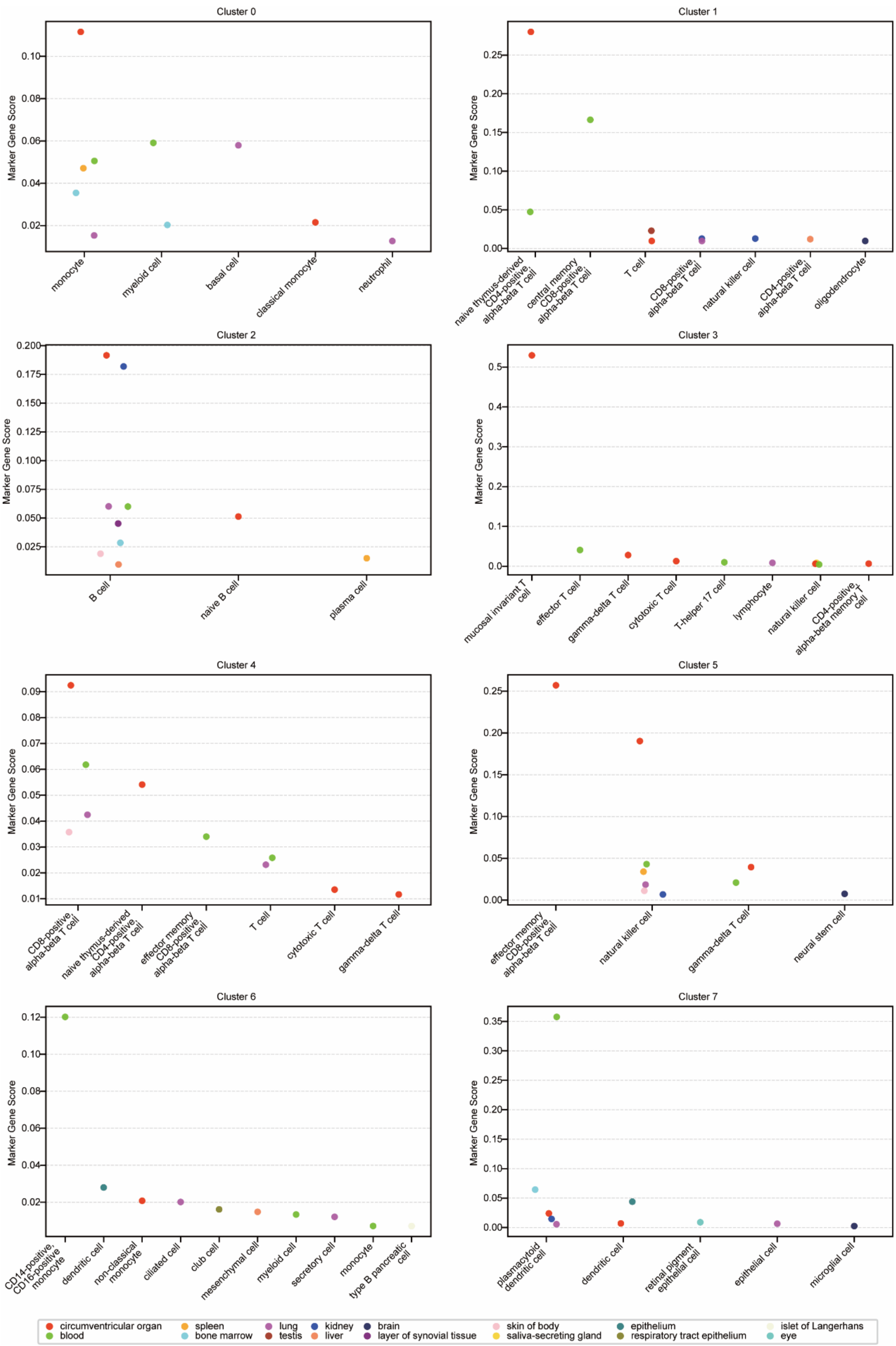


**Supplementary Figure 3. Results of the Cell Identification function for the 10k PBMCs dataset.** Shown are the Marker Gene Scores for each cell cluster. The colors of dots denote different tissues. The x-axis represents cell types, and dots with the same color represent cell types from the same tissue/organ type.

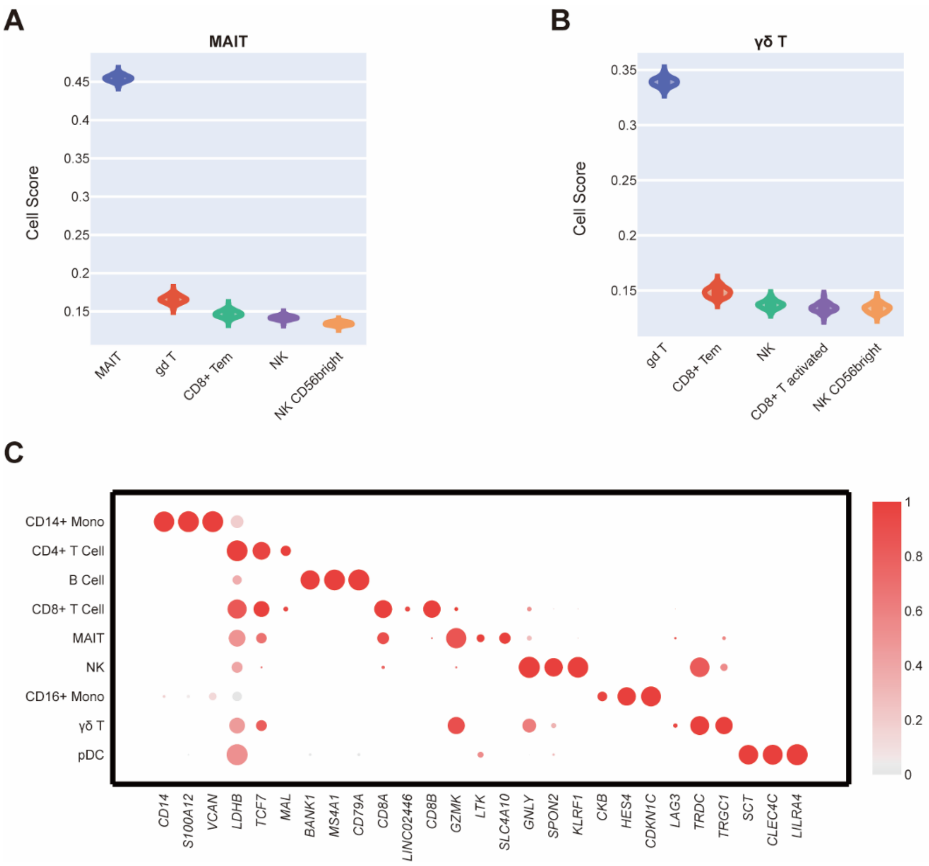


**Supplementary Figure 4. Validation of the "MAIT" and "γδ" cluster annotation results for the 10k PBMCs dataset. (A)** and **(B)** The Reference Comparison function results for Cluster "MAIT" **(A)** and Cluster "γδ" **(B)** by comparing with the PBMCs reference single-cell dataset in the ToppCell database. **(C)** Expression pattern of the top 3 marker genes sorted by COSG score across all annotated cell types.
