## Supplemental Table for "CellClick: an interactive platform for adjustable and accurate cell type annotation in single-cell and spatial omics data"

**Supplementary Table 1 List of cell atlases used in the Reference Comparison function of CellClick**

| Atlas Name | Annotation Category | Publication (Doi) |
|---|---|---|
| Human_Adipose_Subcutaneous | cell_type | 10.1101/2023.09.04.555678 |
| Human_Adipose_Visceral | cell_type | 10.1101/2023.09.04.555678 |
| Human_Breast | broad_cell_type | 10.1038/s41586-023-06252-9 |
| Human_Eye_snRNA | majorclass | 10.1101/2023.11.07.566105 |
| Human_Eye_scRNA | majorclass | 10.1101/2023.11.07.566105 |
| Human_Gut | category | 10.1038/s41586-021-03852-1 |
| Human_Heart | cell_type | 10.1038/s41586-023-06311-1 |
| Human_Blood | major_subset | 10.1016/j.cell.2022.01.012 |
| Human_Kidney | subclass.l1 | 10.1038/s41586-023-05769-3 |
| Human_Liver | author_cell_type | 10.1016/j.cell.2021.12.018 |
| Human_Lung | ann_level_3 | 10.1038/s41591-023-02327-2 |
| Human_Brain_Neuronral | supercluster_term | 10.1126/science.add7046 |
| Human_Brain_None_Neuronal | supercluster_term | 10.1126/science.add7046 |
| Human_Organoid_Neural | annot_level_2 | 10.1038/s41586-024-08172-8 |
| Human_Organoid_Endoderm-Derived | level_2 | 10.1101/2023.11.20.567825 |
| Mouse | cell_type | 10.1093/nar/gkac633 |
